# Density effects on a tropical copepod *Acartia* sp.: implications as live feed in aquaculture

**DOI:** 10.1101/2023.03.02.530601

**Authors:** Hung Quoc Pham, Canh Van Bui, Nam Xuan Doan, Khuong V. Dinh

## Abstract

Calanoid copepod *Acartia* species are major live feeds for the early stages of economically important marine fish in hatcheries in Southeast Asian countries. However, rearing *Acartia* copepods at high densities to increase productivity remains a major challenge. To address the issue, we conducted two experiments on 1) *Acartia* sp. nauplii (1000, 3000, 6000, 9000, 12000, and 15000 individuals L^-1^) and 2) adults (1000, 1500, 2000, and 2500 individuals L^-1^). We assessed key parameters for biomass production: development, survival, and egg production. In general, increased density resulted in longer development time, and lowered survival and egg production, but did not affect the size of adult males and females. Despite survival to adulthood decreasing at higher stocking nauplii densities, the number of surviving adults was highest at a stocking density of 12000 ind L^-1^. Egg production was very low which may be the result of high egg predation. The total eggs harvested were highest at the lowest adult density. These results are essential for the biomass production of *Acartia* sp. in central Vietnam.

## Introduction

Copepod biomass production has become one of the major interests in aquaculture for the last two decades (Drillet *et al*. 2006, Doan *et al*. 2018, Grønning *et al*. 2019, Nguyen *et al*. 2020, Drillet *et al*. 2011) due to the high nutritional value, size variations and behaviour of copepods which may be relevant for larvae of many aquaculture species (Drillet et al. 2011, Drillet et al. 2006). Calanoid copepod species have high levels of highly unsaturated fatty acids (HUFA) (Drillet *et al*. 2008, Drillet et al. 2006, Rayner *et al*. 2017b, Rasdi and Qin 2016). Tropical calanoid copepods are ideal live feeds for important aquaculture fish larvae including grouper *Epinephelus* sp. and cobia *Rachycentron canadum* (Lee et al., 2010), snapper *Lutjanus guttatus* (Burbano et al., 2020). In tropical and subtropical aquaculture ponds, copepod density is typically lower than 400 adult individuals L^-1^ (Blanda *et al*. 2015, Grønning et al. 2019) which may result in low production (Grønning et al. 2019).

Culturing calanoid copepods at high densities remains a great challenge (reviewed in Drillet et al., 2011). Indeed, the negative effect of density on the ovigerous rate of *Pseudodiaptomus annandalei*, a tropical calanoid copepod was observed at > 400 individual L^-1^ (Rayner *et al*. 2017a). For *Acartia* species, the density effect on the copepod performance and production has been tested at much higher densities, from 125 to 8,000 adults L^-1^ for *Acartia bilobata* (Chintada *et al*. 2021) and from 5,000 to 45,000 adults L^-1^ for *A. tonsa* (Torres *et al*. 2022). The challenge is to meet a large number of copepods raised in artificial conditions for marine fish larvae rearing on a commercial scale.

In this study, we assessed the performance of a tropical calanoid copepod *Acartia* sp. at different nauplii and adult densities. We determined the development time, survival, size of adult males and females and egg production. These parameters are essential for the biomass production of copepods for aquaculture purposes.

## Materials and methods

### *Copepod Acartia* sp

Copepod *Acartia* sp. were collected from aquaculture ponds in Cam Ranh area. This could be a new *Acartia* species (D.H.Q Vu et al., in prep). Copepods were kept at Cam Ranh Aquaculture Experiment Center, Institute of Aquaculture, Nha Trang University. Copepods were reared in water with a salinity of 30 ppt, room temperature 28-31°C, natural light, and gentle aerations. *Acartia* sp. were fed with *Isochrysis* algae at a concentration of ∼40,000 cells/mL, 3 times a day at 6h, 14h, and 22h. The water in the culture tanks was daily exchanged by 20 -30% by a siphoned tube with a 25 µm mesh end to avoid losing copepod. *Acartia* sp. To collect *Acartia* sp. for the experiments, copepods were sucked from the culture tanks with siphon tubes through a 200 µm filter to retain the adults.

### Water source

Seawater is taken from Canh Ranh Bay into a pond containing 3000 m^2^ with a salinity of 35 ppt, then it will be pumped through a 5 µm filter bag into two 4 m^3^ composite tanks, treated with 100 ppm chlorine. After chlorine treatment, all residue of chlorine was removed by aerations and water was pumped through a 0.5 µm filter cylinder system into 200L indoor tanks for use.

### Microalgae

The feed used for experimenting on this copepod was *Isochrysis galbana*. Algae were grown under laboratory conditions at 26°C with a semi-continuous culture method. Algae are grown in a 5,000 mL triangle with a 24/24 hour light using 4 60W LEDs (1.2 m long). The medium used for algae culture was f/2 medium. The water used for algae culture has a common source with the copepod culture water. Before use for algal culture water was boiled and left to the same temperature as the culture and has a salinity of 25 ppt.

#### *Experiment 1: Determining the effects of stocking density of Acartia* sp. nauplii

To produce the nauplii for the experiment, adults *Acartia* sp. were collected from the ponds and transferred to a 200-L tank containing 180 L of treated seawater at a density of 1,000 individuals L^-1^. Subsequently, the algae *Isochrysis* was added at a density of 40,000 cells/mL. Egg production of *Acartia* sp. was monitored and collected every 12 hours. In details, eggs on the bottom of the tank were collected by siphoning, and the water containing eggs was filtered through a 50 µm filter, the eggs were stored at 4°C. After 4 times of egg collection, the eggs were hatched in 25 ppt seawater at room temperature 28-30°C. After 48 hours of incubation, nauplii were collected using a net (mesh size = 25 µm) and quantified for the experiment.

Nauplii were transferred into 5-L plastic bottles with experimental densities with the following treatments: Treatment 1: 1,000 nauplii L^-1^; Treatment 2 3,000 nauplii L^-1^; Treatment 3 6,000 nauplii L^-1^; Treatment 4 9,000 nauplii L^-1^; Treatment 5 12,000 nauplii L^-1^; Treatment 6 15,000 nauplii L^-1^. Each treatment had 3 replications. We daily took 50 ml of samples from each of the experimental bottles to determine the percentage of nauplii, copepodites, and adults in each sample. Adult males and females were collected from the experimental bottles and the size of the prosome was measured using a stereomicroscope (Truong *et al*. 2020, Dinh *et al*. 2021). To determine the survival, we took a subsample (250 ml) from each of the experimental bottles on day 10 and counted the number of adults to estimate the survival from nauplii to adulthood.

#### *Experiment 2: Determination of optimal stocking density of Acartia* sp. *adults*

To have the adult copepods, we collected nauplii from 6 × 200-L rearing tanks. Copepods were fed daily with *Isochrysis galbana* algae at a density 40,000 cells mL^-1^, 2 times a day, and with a water exchange rate of 20% volume per day (Doan et al. 2018). After 8 days, adult copepods were collected using a 150 µm filter.

Adult copepods were assigned to 20 L tanks containing 5 L of filtered seawater. Density in stocking treatments was Treatment 1 1,000 individuals L^-1^; Treatment 2 1,500 individuals L^-1^; Treatment 3 2,000 individuals L^-1^; Treatment 4 2,500 individuals L^-1^. Each treatment was repeated 5 times. Egg production was monitored by collecting eggs in each of the experimental bottles every other day. Eggs were concentrated in 100 ml seawater, and gently mixed. We collected a subsample of 1 ml and the number of eggs in the subsamples was counted. Subsequently, the number of eggs produced from each of the experimental bottles was estimated from days 2 to day 14. We also calculated the total egg production per treatment and the average number of eggs produced by a female on day 14 of the experimental period.

### Data analysis

The data is processed and displayed graphically on Excel software. Using SPSS 20v software to analyze one-way variance (one-way ANOVA) with the Duncan test. Data are expressed or visualized in graphs as mean ± standard error (Mean ± SE).

## Results and Discussion

### Experiment 1: Determination of optimal culture density of *Acartia* sp. nauplii

#### Development time at different densities

The development time to reach 100% maturity from the nauplii stage varied by culture densities (Figure 1). The development time was also longer at higher experimental densities. Initially, the nauplii phase lasted only 4 days at experimental densities below 9000 individuals L^-1^. While it was 5 days at the density of 12000 individuals L^-1^ and 15000 individuals L^-1^. In the copepodite stages, the density effect on the copepodite development was more pronounced compared to the nauplii stages. Indeed, at a density of 100 individuals L^-1^, the copepodite stages lasted only 2 days, but it was 6 days when the experimental density increased to 12,000 individuals L^-1^. As a result, the population of *Acartia* sp. completed maturation and also varied between experimental densities. Specifically, at the stocking density of 100 individuals L^-1^, it only took 6 days for the population to reach full maturity, but at the density of 12,000 individuals/L and 15,000 individuals L^-1^, the development time was 10 days. At the densities of 3,000, 6,000, and 9,000 individuals L^-1^, the development time was 7, 8 and 9 days, respectively.

**Figure 1.**
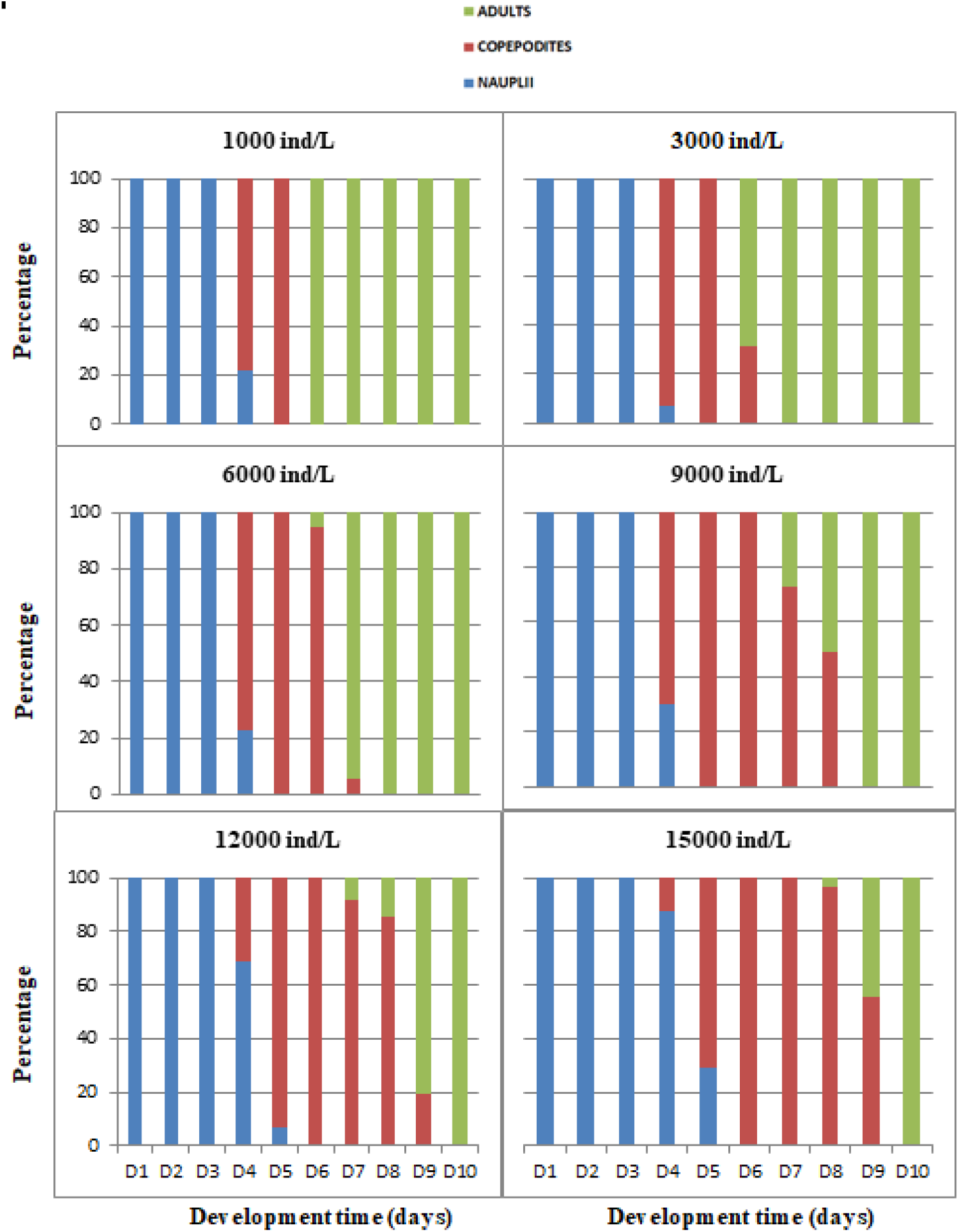
Development time (day) at different densities.

**Figure 1.**
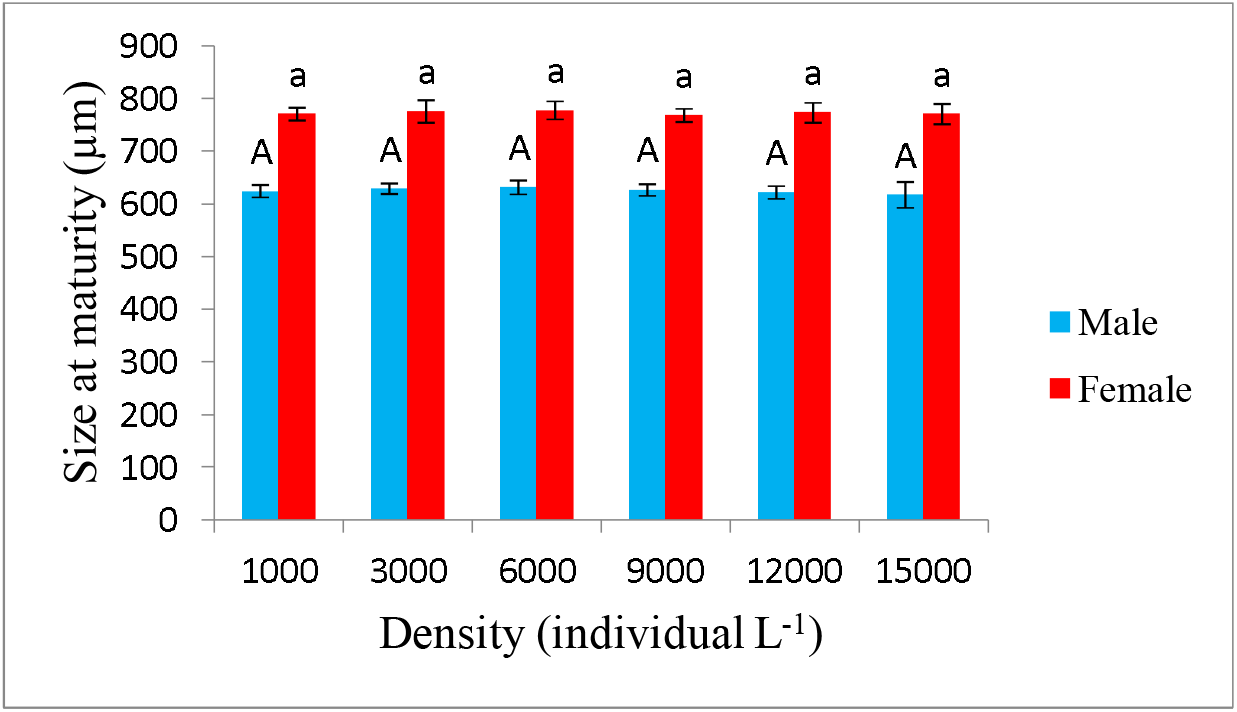
The size of males and females at different densities.

#### The size of adults at different densities

The size of both males and females was not affected by the culture density. There was no statistically significant difference (*p* > 0.05) between different experimental densities. Male size ranged between 617 -631, while the female ranged between 768 – 778 µm (Figure 2).

#### Survival from nauplii to adult stage

The survival from nauplii to the adult stage was shown in Figure 3 A. Overall, the survival was lower at higher densities. At the densities of 1000-3000 individuals L^-1^ survival was highest. At a density of 15,000 individuals L^-1^ the survival was lowest, 39.5% surviving to adulthood. At densities of 6,000, 9,000, and 12,000 individuals L^-1^ survival was not statistically different from each other (p > 0.05). Despite the lower survival percentage from nauplii to adulthood at higher densities, the number of individuals who reached the adult stage was higher with increased stocking density (Figure 3 B), until a stocking density of 12,000 individuals L^-1^, which had the highest number of adults. Increasing the stocking density from 12,000 to 15,000 individuals L^-1^ did not result in a higher, but a lower final density of adults (Figure 3B), which could be linked to very high mortality (Figure 3B).

**Figure 3.**
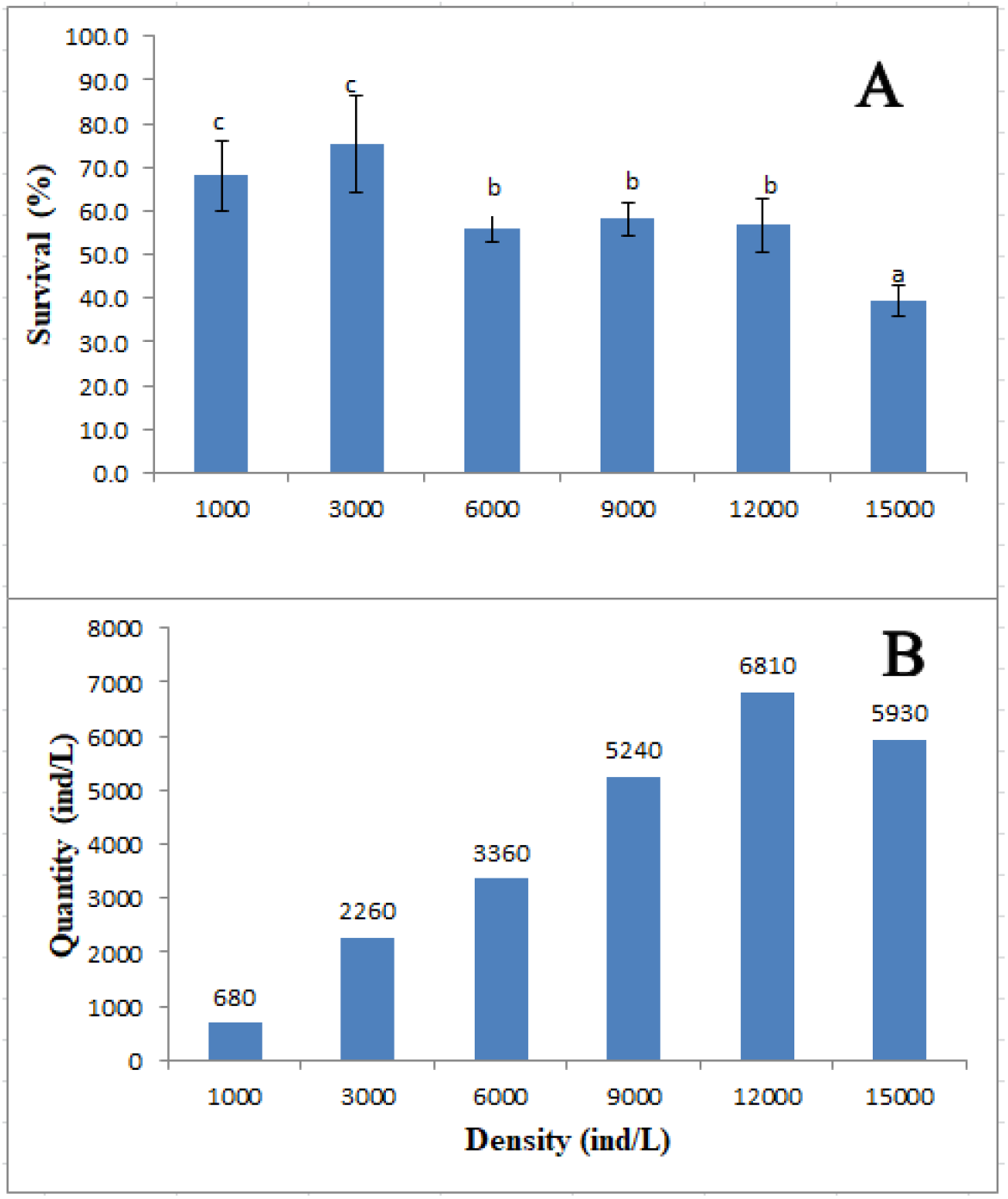
Survival (A) and the number of individuals reached the adult stage (B) at different densities.

**Figure 4.**
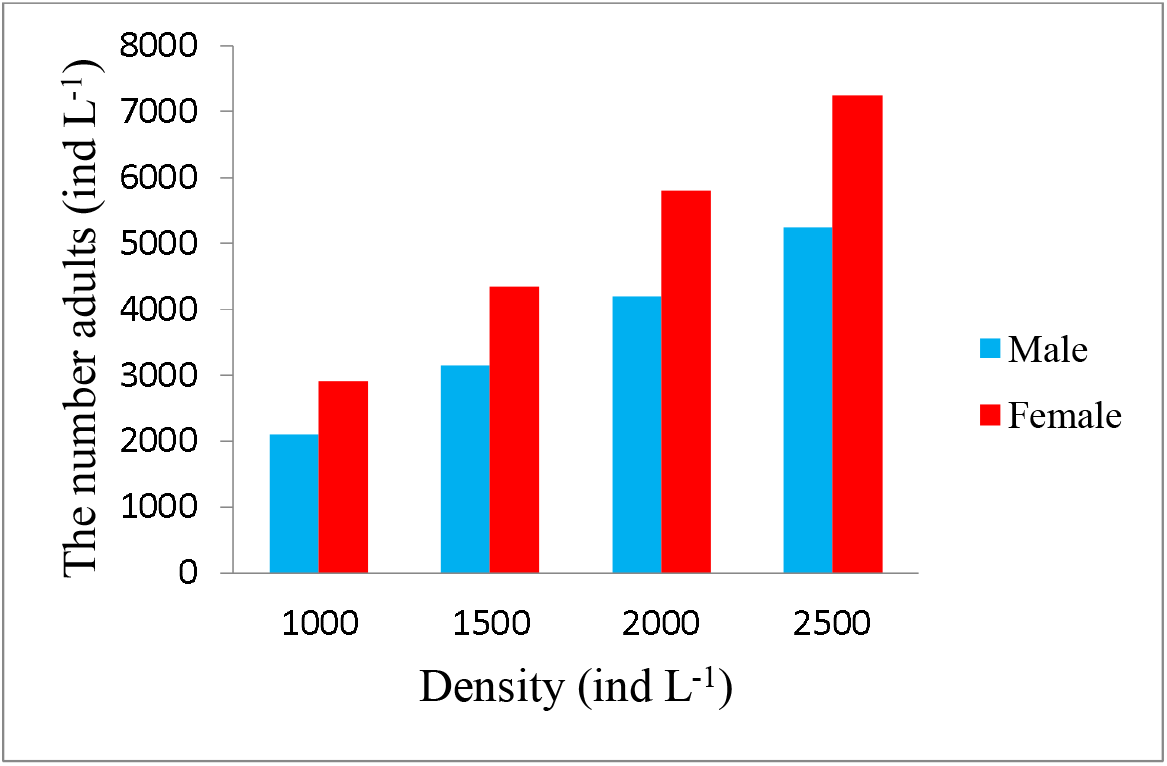
Number of males and females at different densities.

### Experiment 2: Effects of adult stocking density

#### The number of males and females at different densities

*Acartia* sp. with an initial sex ratio was 42% and 58% females. We could convert the number of individuals corresponding to each experimental density.

#### Effect of stocking density on the egg production

The egg production of *Acartia* sp. at experimental densities tended to increase from day 2 to day 8 and then gradually decreased to day 14 (Figure 5). The number of eggs was high from day 6 to day 10. When studying the effect of density at the beginning to the spawning of *Acartia sinjiensis*, the total number of eggs obtained daily was high in the first 3 days and gradually decreased with experimental density, the average number of eggs did not differ over time (Camus and Zeng 2009).

**Figure 5.**
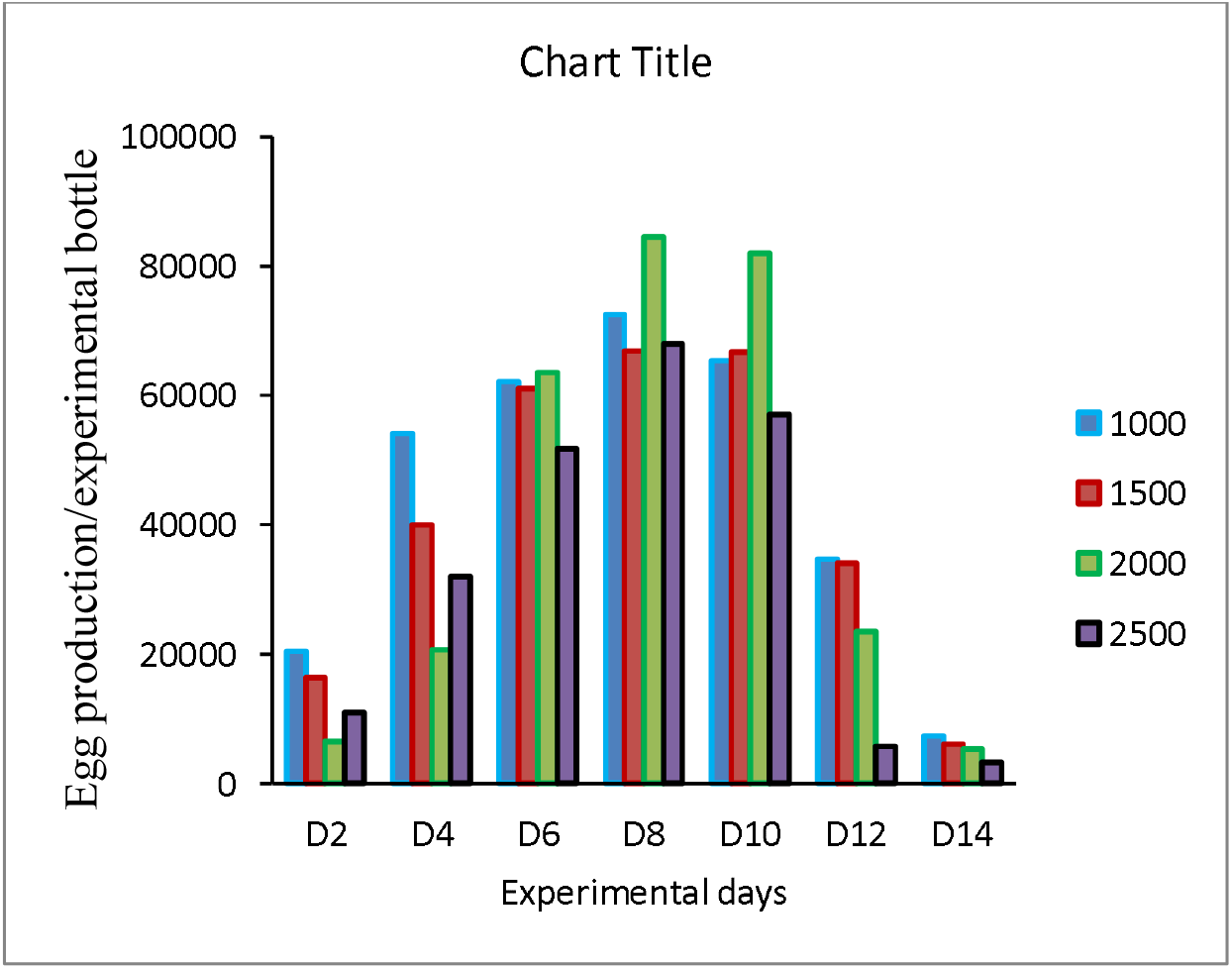
Egg production at different densities over the experimental period.

#### Effect of stocking density on total eggs harvested

Figure 6 shows the total number of eggs obtained at different experimental densities. The number of eggs harvested at the experimental density of 1000 individuals L^-1^ was the highest with an average of 31,6739 eggs. The experimental density of 2,500 individuals L^-1^ had the lowest total number of eggs with an average number of 23,5024 eggs. Egg production also tended to decrease with an increasing stocking density of *Acartia tonsa* from 100 to 2500 individuals L^-1^ (Franco *et al*. 2017). When assessing the effect of adult density on the fertility of *Acartia tropica*, the author also noted that a density of 1,000 individuals/L is suitable for egg and nauplii culture to be used as live food (Wilson *et al*. 2022). A density of 1000 adults L^-1^ is also recommended for rearing *Acartia bilobata* (Chintada et al. 2021) to achieve optimal total egg production.

**Figure 6.**
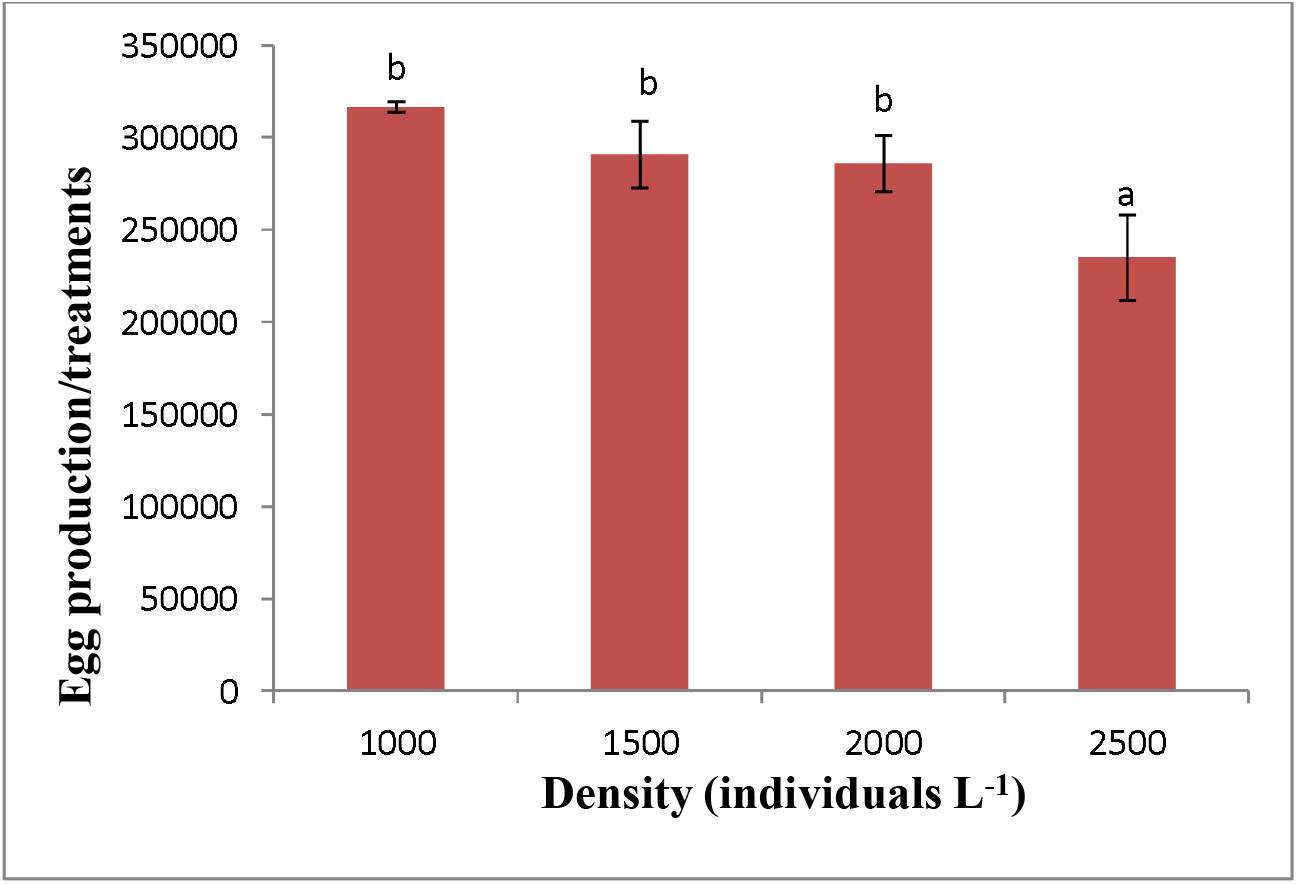
Total eggs obtained from 5 replications of a treatment at different densities.

**Figure 7.**
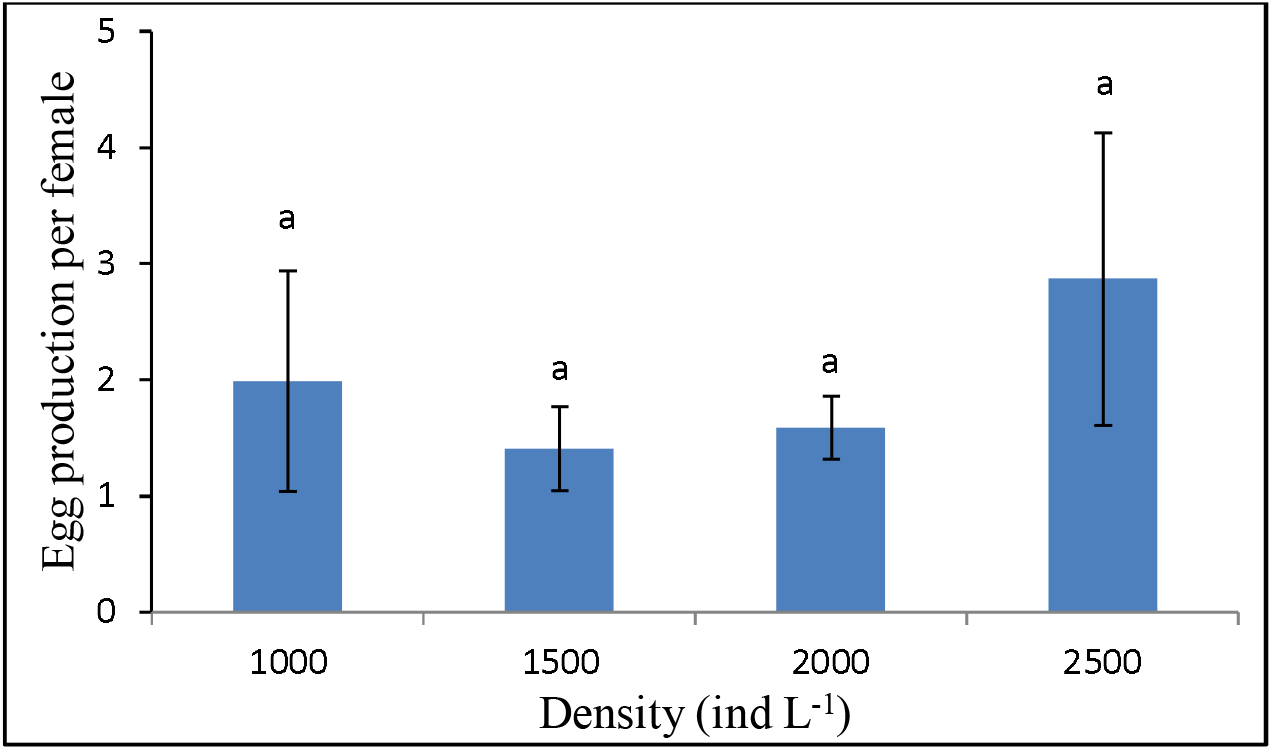
Fertility of *Acartia* sp. one on the day 14 at different densities.

**Figure 8.**
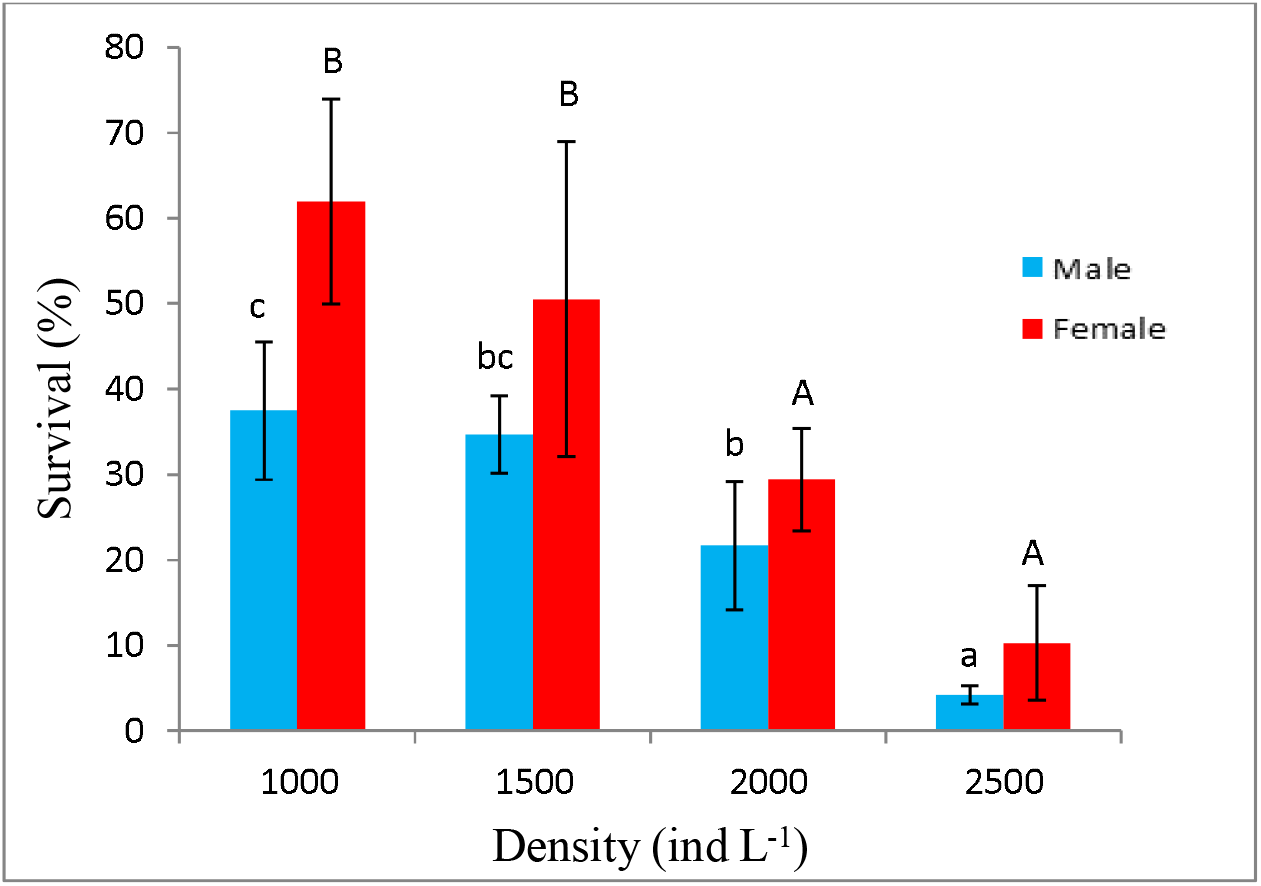
Survival after 14 days of culture at different densities.

##### Effect of density on the average number of eggs produced on the day 14th

The egg production of *Acartia* sp. was generally low (< 3 eggs/female). The average number of eggs per female after the experiment was very low compared to *Acartia tonsa* (e.g., 7 -43 eggs per female/day, Torres et al. 2022, Krause *et al*. 2017). The difference in egg production of *Acartia* sp. could be explained by the species difference, or the food for copepods e.g. *Isochrysis galbana* in this study, but *Rhodomonas baltica* in other studies (Torres et al. 2022, Krause et al. 2017). Finally, *Acartia* copepods may feed on their eggs (egg predation), particularly at high densities (Vu *et al*. 2017). An egg collector system is required to allow the egg to fall quickly into the bottom of the experimental containers, thereby avoiding egg predation by adults (2022).

#### Effect of density on survival

In general, the survival rate after 14 days of the experiment decreased with the increase in stocking density. Specifically, the survival rate in both males and females was highest at the density of 1000 individuals L^-1^ and there was no statistically significant difference (*p* > 0.05) compared with the density of 1,500 individuals/L, and density of 2,500 individuals/L has the lowest survival rate. When studying the stocking density of *Acartia sinjiensis*, the survival rate after 12 days of culture only fluctuated 10-12% at a culture density of 1,000-2,000 individuals L^- 1^ (Camus and Zeng 2009). But in this experiment, the survival rate of *Acartia* sp. fluctuated 20-60% at the density of 1,000-2,000 adult individuals L^-1^. Increased stocking density of adult *Acartia tonsa* also resulted in a significant decrease in survival (Franco et al. 2017). The negative density effect on the survival of *Acartia* species has been observed in *A. bilobata* (Chintada et al. 2021), which could be a result of increasing competition for food and space (Chintada et al. 2021, Franco et al. 2017, Sibly *et al*. 2000). In the same culture density, the survival of *Acartia* sp. males was lower than *Acartia* sp. Females, which can be linked to the generally higher sensitivity of males to stressors caused by higher oxidative damages (Rodríguez-Graña *et al*. 2010) and a shorter lifespan than females in calanoid copepods (Truong et al. 2020, Rodríguez-Graña et al. 2010).

### Conclusions and recommendations

Stocking density of nauplii *Acartia* sp. different baselines had a marked effect on population growth as well as survival to adulthood. From the above research results, the initial nauplii density was 3,000 nauplii L^-1^, which is suitable for stocking the copepod *Acartia* sp. Density of individuals *Acartia* sp. The initial maturation also has an impact on the reproductive health of the population, and the density of 1000 individuals L^-1^ is suitable for rearing *Acartia* sp. egg harvest.

The eggs of *Acartia* copepods have many different forms corresponding to different hatching times such as diapause eggs, and resting eggs. The ratio of each type depends on each condition, and there is a density factor in it. Therefore, it is necessary to study and evaluate the hatching rate of eggs when reared at different densities.

## Data availability

Data are available at Nha Trang University research server.

### Ethics statement

Experimental work on copepods does not require ethical approval or a license for working with animals.

## Acknowledgments

This research was funded by Nha Trang University, Vietnam with the grant code TR2021-13-16.

